# Androgen-*dmrt1* positive feedback programs the rice field eel (*Monopterus albus*) sex transdifferentiation

**DOI:** 10.1101/595306

**Authors:** Bin Wen, Xiancheng Qu, Lisha Pan, Jianzhong Gao, Haowei Wu, Qian Wang

## Abstract

The rice field eel *Monopterus albus* is a hermaphroditic protogynous fish species that undergoes sex reversal from female to male. However, the potential mechanisms underlying the process of sex transformation are still unclear. We analyzed and compared the gene sequence of *M. albus dmrt1* 5′ upstream region and its potential transcription factor binding sites with other known species and examined the *in vitro* effects of testosterone (T) on the expression levels of *dmrt1a* and *foxl2* in the ovotestis. Moreover, we cloned and analyzed the expression of genes encoding enzymes, 11β-hydroxylase (*11β-h*) and 11β-hydroxysteroid dehydrogenase (*11β-hsd*), involved in the production of 11-ketotestosterone (11-KT). The results showed that, compared with other fish species, *M. albus dmrt1* 5′ upstream region contained unique androgen response elements (AREs) with one on the sense strand and the other one on the antisense strand, indicating a crucial role for androgens in the transcriptional regulation of *dmrt1*. The expression of *dmrt1a* was induced but the expression of *foxl2* was inhibited by T manipulation *in vitro*, suggesting that blood androgen could activate the transcription of *dmrt1* in the ovotestis. Moreover, the expression levels of *11β-h* and *11β-hsd2* were predominantly expressed in testis, much less in ovotestis, and barely in ovary, suggesting the production of 11-KT during sex reversal. Androgens are synthesized in large amounts during sex reversal, leading to the promotion of *dmrt1* transcription, and thus, gonadal somatic cells transdifferetiation. Overall, androgen-*dmrt1* positive feedback programs the *M. albus* sex reversal.

## Introduction

Androgens in teleosts are essential for inducing male phenotype and male gametogenesis, and female-to-male sex reversal in some species. Both testosterone (T) and 11-ketotestosterone (11-KT) are detected in males, the latter being the potent androgen responsible for testicular development (1). The regulation of enzymes involved in the biosynthesis of 11-KT are critical for teleostean reproduction. 11β-hydroxylase (*11β-h*) and 11β-hydroxysteroid dehydrogenase (*11β-hsd*) are two important steroidogenic substrates for the production of 11-KT (2, 3). During spermatogenesis substantial changes in the expression level of *11β-h* are observed in rainbow trout *Oncorhynchus mykiss* (4, 5), medaka *Oryzias latipes* (6), Atlantic salmon (7), Nile tilapia *Oreochromis niloticus* (8, 9) and catfish *Clarias batrachus* (10). Similarly, *11β-hsd* transcripts are present in the steroidogenic tissue of *O. mykiss* and its transcriptional signals were observed in the Leydig cells of testes, in the thecal cells of early vitellogenic ovarian follicles, and in the thecal and granulosa cells of midvitellogenic and postovulatory follicles (2). Also, *11β-hsd2* is expressed in various tissues of *O. niloticus*, with the highest expression level observed in the testis (3).

Many genes are known to be involved in gonadal differentiation in vertebrates. *Dmrt1*, a gene that encodes a transcription factor with a DM-domain, is one of the essential genes controlling testicular differentiation in mammals, birds, reptiles, amphibians and fish (11-13). In *O. mykiss*, for example, *dmrt1* is expressed during testicular differentiation but not during ovarian differentiation (14). *Dmy* is enough for male development in *O. latipes* and loss of *dmy* in XY medaka causes male-to-female sex reversal (15-17). *O. latipes* also has an autosomal copy of *dmrt1* which is expressed in testis later than *dmy* but is essential for testis development (18, 19). *Dmrt1* is not only associated with testis development, but also, may be crucial for the ovary differentiation in zebrafish (20); however, Webster et al. (21) reported that *dmrt1* is dispensable for ovary development but necessary for testis development by regulating *amh* and *foxl2*. Wen et al. (22) observed that *dmrt1* expression was 70 times higher in the testis of olive flounder *Paralichthys olivaceus* than in the ovary. Also, in European sea bass, the expression of *dmrt1* is increased in testis but decreased in ovary (23).

The rice field eel *Monopterus albus* is a hermaphroditic fish species that undergoes sexual reversal from a functional female to a male (24). The *M. albus* is emerging as a specific model for studying vertebrate sexual development due to its small genome size and naturally occurring sex reversal (25). Yeung et al. (26, 27) examined the effects of exogenous androgens on sex reversal and sex steroid profiles in the female of *M. albus*. He et al. (28) observed the ovarian differentiation, morphogenesis and expression of some gonadal development-related genes in *M. albus*. Several genes related to sex determination and differentiation have been identified in *M. albus*, including *cyp19a1a* (29), *sox9a* (30), *cyp17* (31), *sox17* (32), *dmrt1* (33), *jnk1* (34), *foxl2* (35), miRNAs (36) and gonadal soma-derived factor (37). We also investigated the transcription profiles of some genes involved in gonad development and sex reversal in the *M. albus* (38, 39). Moreover, a chromosome-scale assembly of *M. albus* genome is currently available (40). However, the biology events and potential mechanisms underlying the process of female-to-male sex reversal in this species are still unclear.

Huang et al. (33) reported that not only is *dmrt1* expressed specifically in the gonads of *M. albus*, but its multiple isoforms are differentially co-expressed during gonad transformation. Also, Sheng et al. (41) observed that the *dm* genes are involved in the sexual differentiation of *M. albus*. However, the regulation of *dmrt1* in *M. albus* during sex reversal remains largely unknown. As an important *cis*-acting element, core promoter plays pivotal role in the regulation of metazoan gene expression (42, 43). We assumed that a) the 5′ flanking region of *M. albus dmrt1* contains unique promoter motifs that regulates its transcription during sex reversal, b) there is no sex determination gene in *M. albus*, which sex transformation is an evolution process, c) and thus, the process of female-to-male sex reversal is controlled by endocrine regulation and sex hormones play a vital role during this process. To test this hypothesis, we analyzed and compared the gene sequence of *M. albus dmrt1* 5′ upstream region and its potential transcription factor binding sites with other fish species and examined the *in vitro* effects of T on the expression of *dmrt1a* and *flox2* in the ovotestis. Moreover, we cloned and examined the expression patterns of genes encoding enzymes, *11β-h* and *11β-hsd2*, involved in the production of 11-KT in the testis, ovotestis and ovary tissue, so as to reveal the molecular mechanism of sex reversal in *M. albus*.

## Results

### Nucleotide sequence of dmrt1 5’ upstream region

The 5′ flanking region of *M. albus dmrt1* was 1421 bp in size. *In silico* functional analysis showed the transcription binding sites for AP-1, Oct-1, Zen-1, USF, C/EBPa, GATAx, STATx, Foxd3, SRY, Dmrt3, Ftz, ERE, ARE and Sox family of transcription factors (Fig. 1). Specifically, in comparison with other known fish species, the sequence of *M. albus dmrt1* 5′ upstream region contained two unique androgen response elements (AREs), with one on the sense strand (−638 bp ∼ −648 bp) and the other one on the antisense strand (−903 bp ∼ −917 bp) (Supplementary Table 2).

**Fig. 1.**
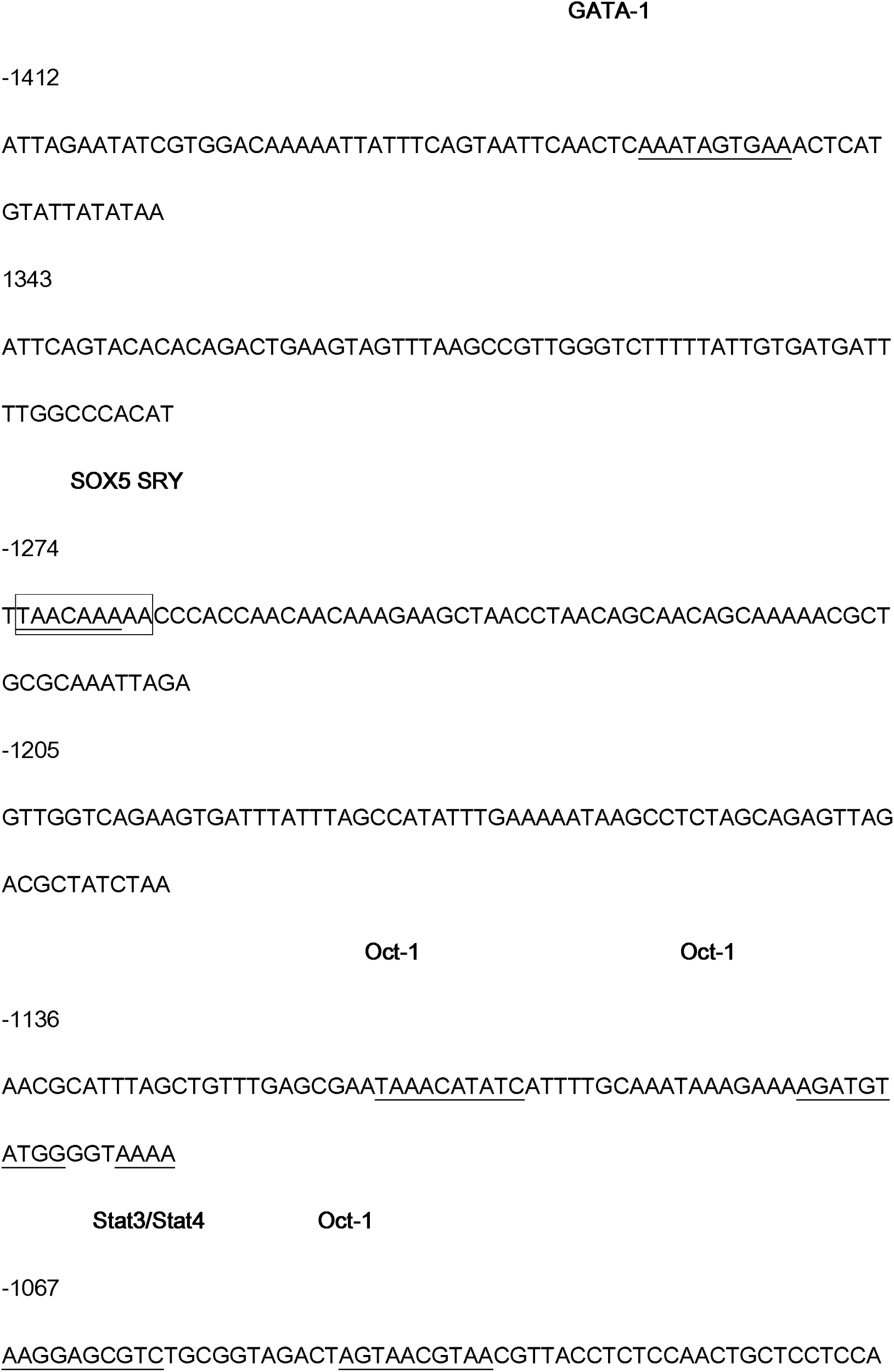
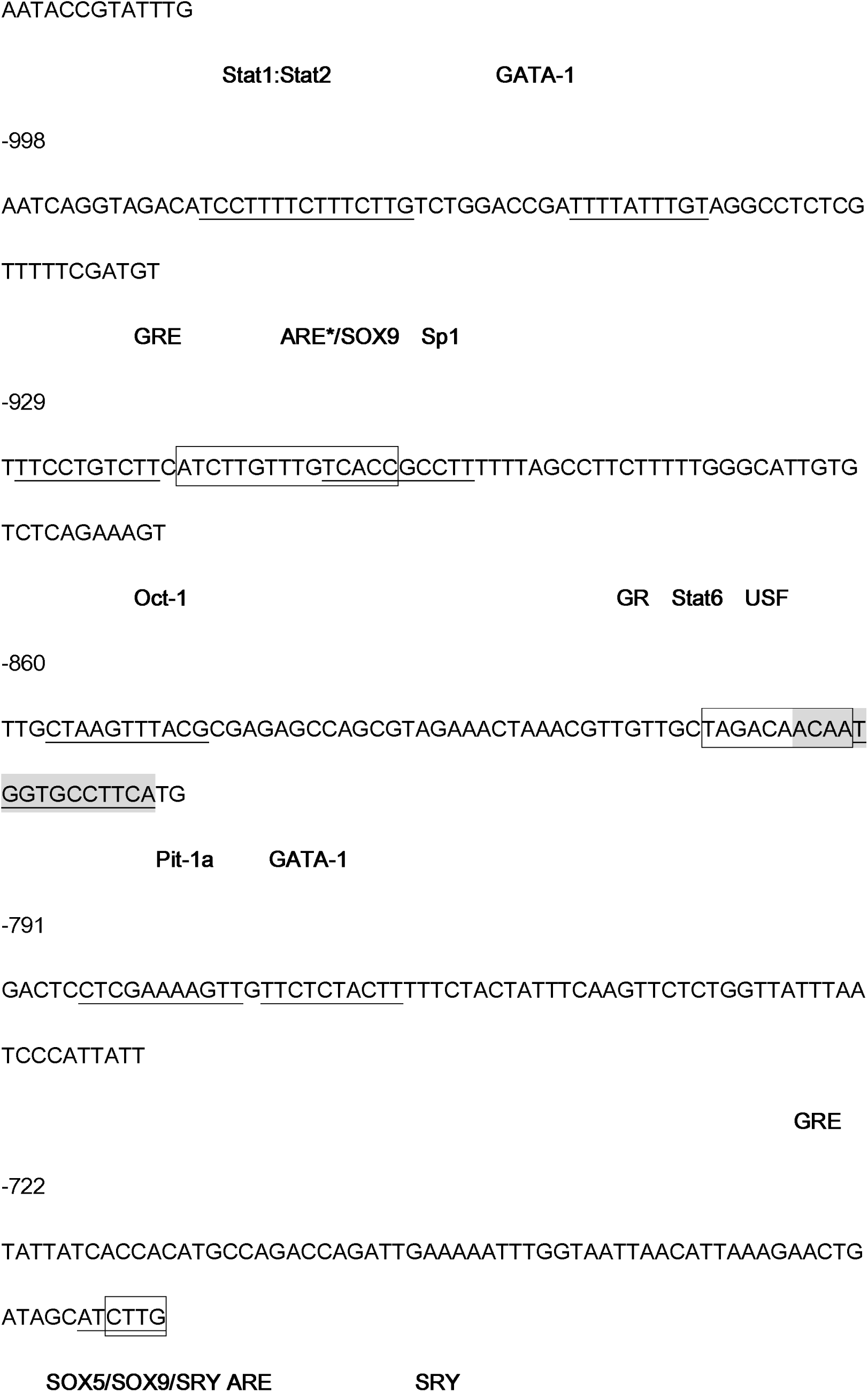
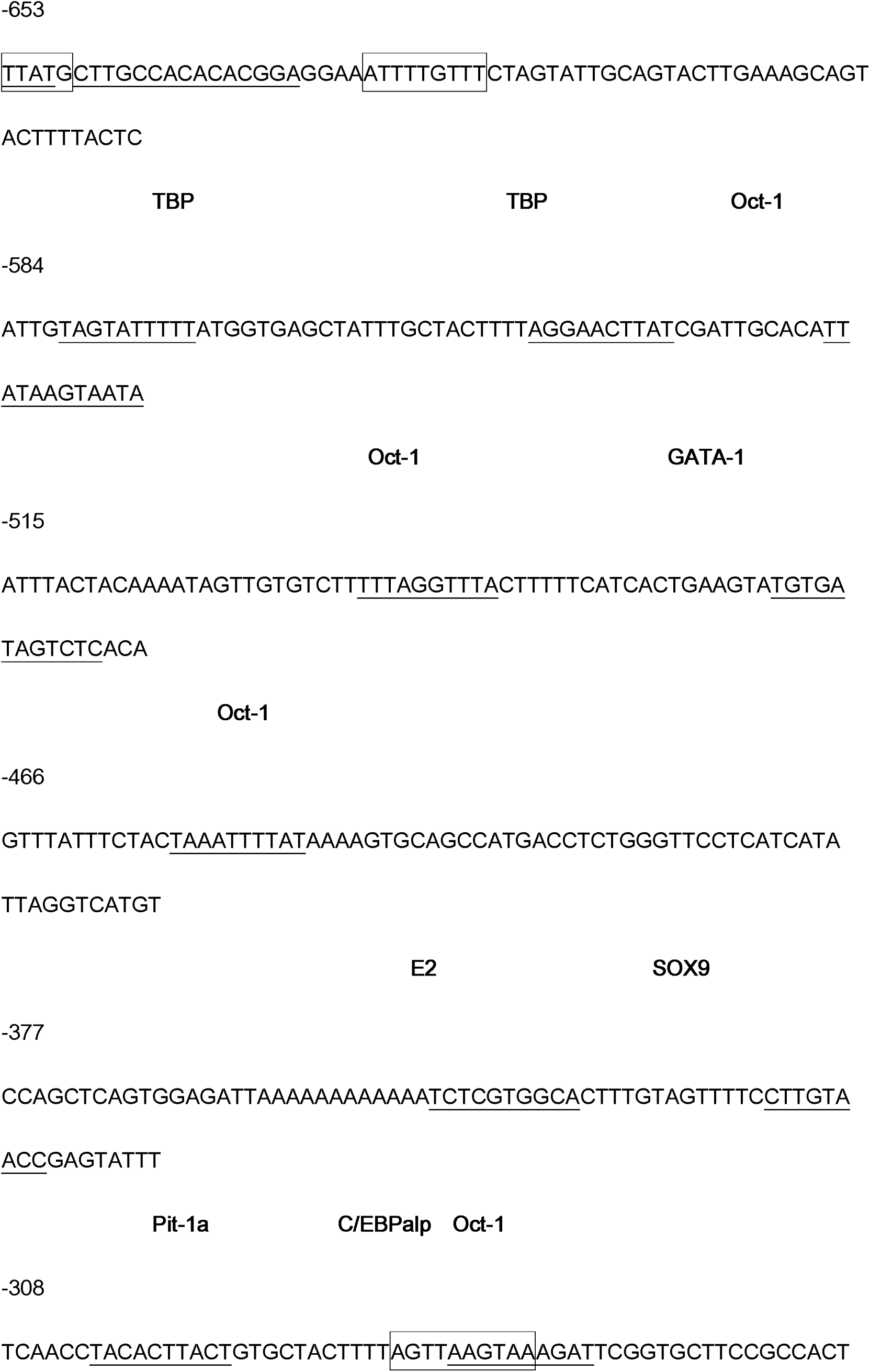
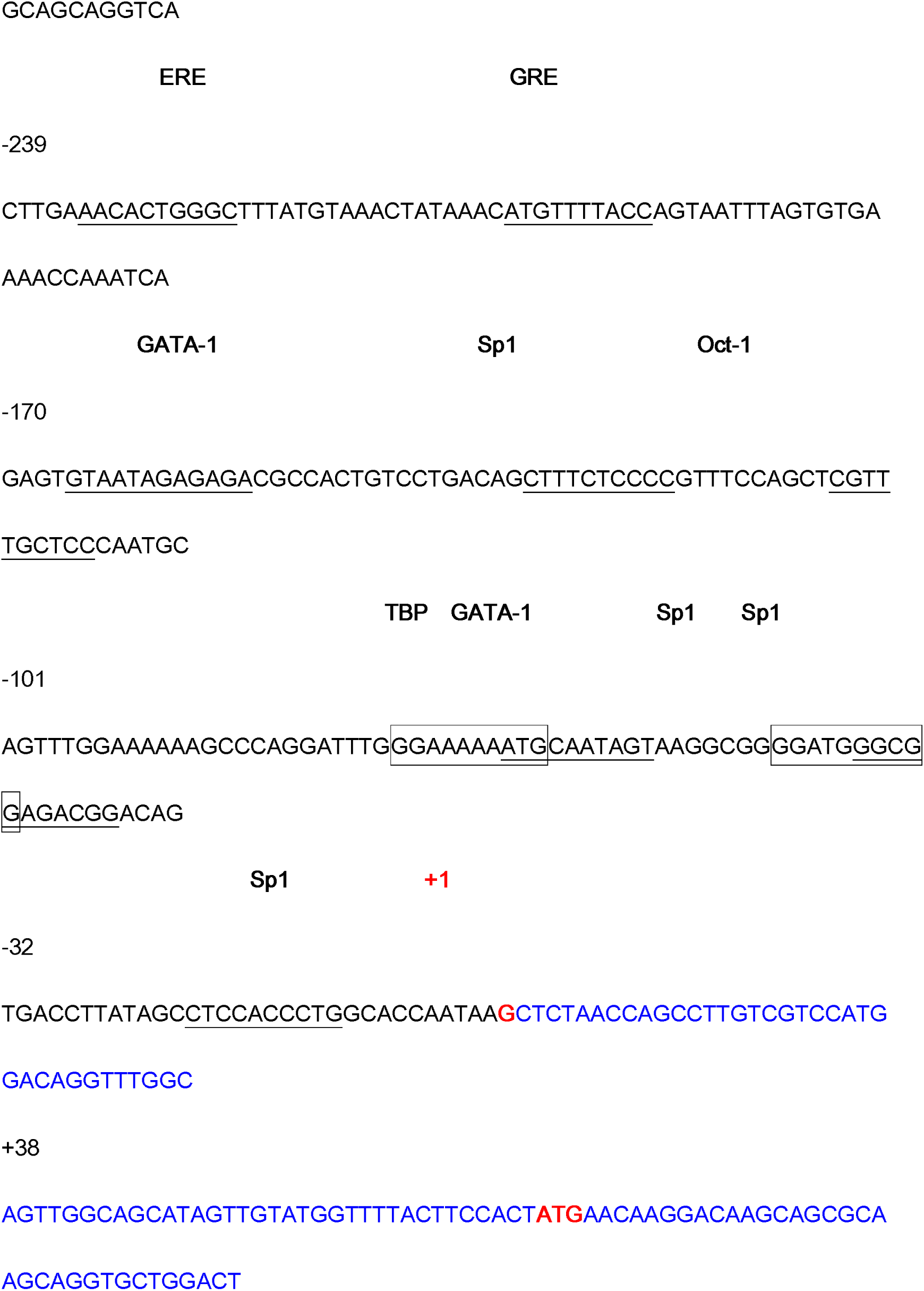
Nucleotide sequence of *M. albus dmrt1* 5′ upstream region with its potential transcription factor binding sites. The potential transcription binding sites are boxed or underlined. (*) indicates ARE on the antisense strand. (+1) means transcription start site.

### Histological change

After 6 hours of tissue culture, cells began to migrate from the periphery of the gonad. Growing tissue appeared after about 5-6 days and the cells were closely arranged and gradually sparse around tissue. There were three types of cells including spindle-shaped fibroblasts, elliptical nuclei; polygonal epithelioid cells; round germinal stem cells, mononuclear or multicellular. The number of cells increased dramatically forming a single layer within 5-6 days. The epithelioid cells and germinal stem cells began to vacuolate and were gradually apoptosis with the extension of culture time, and fibroblasts dominated after 11-12 days (Fig. 2A-D).

**Fig. 2.**
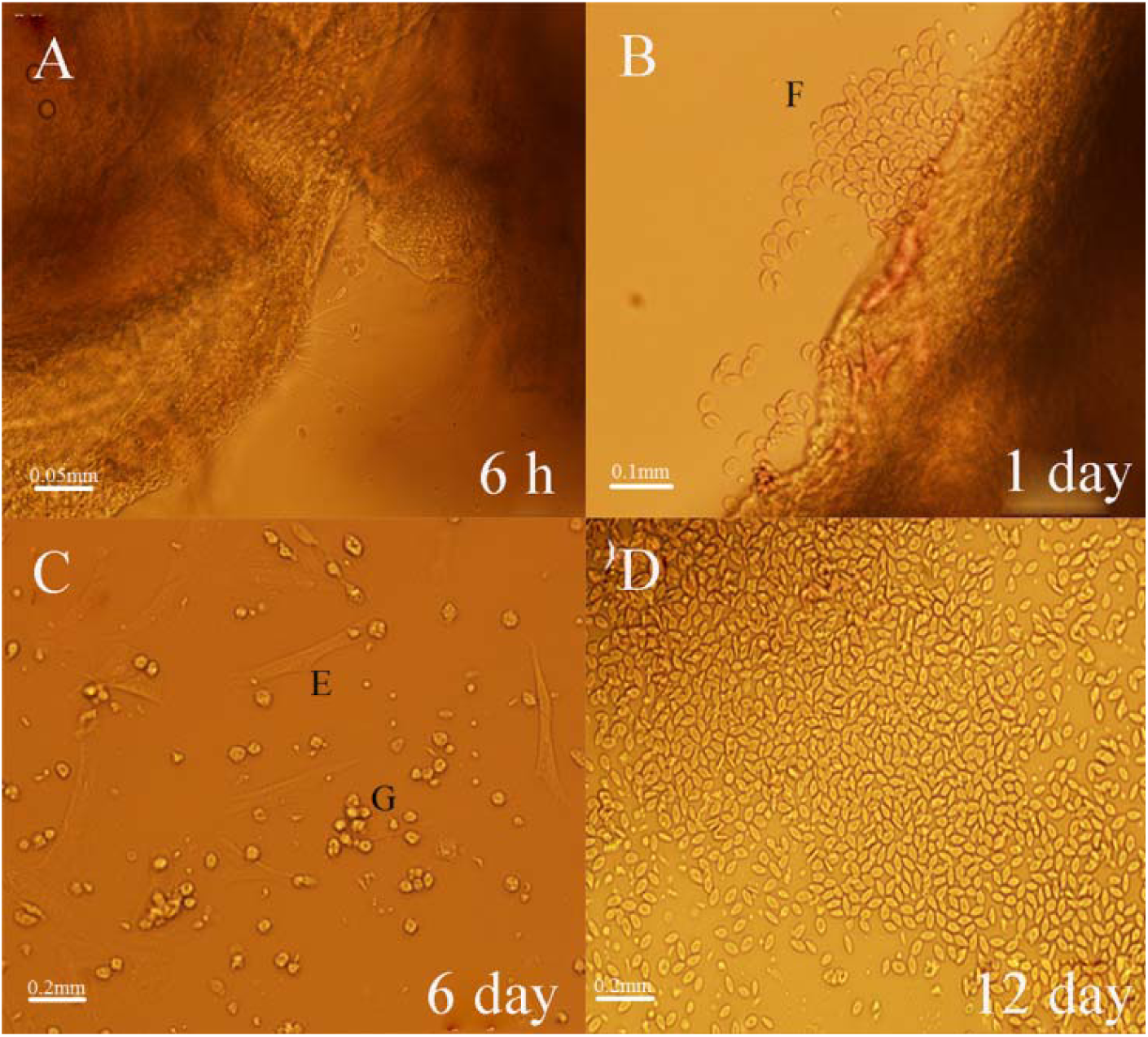
*In vitro* culture of ovotestis in *M. albus*. A: tissue culture after 6 hours; B: tissue culture after 1 day; C: tissue culture after 6 days; D: tissue culture after 12 days. E: epithelioid cells; F: fibroblast cells; G: germinal stem cells.

### Effects of T on the expression levels of dmrt1a and foxl2

On day 6 and day 12, with increasing concentrations of T, the expression level of *foxl2* was significantly decreased (*p* < 0.05) (Fig. 3A) but the expression level of *dmrt1a* was significantly increased (*p* < 0.05) (Fig. 3B).

**Fig. 3.**
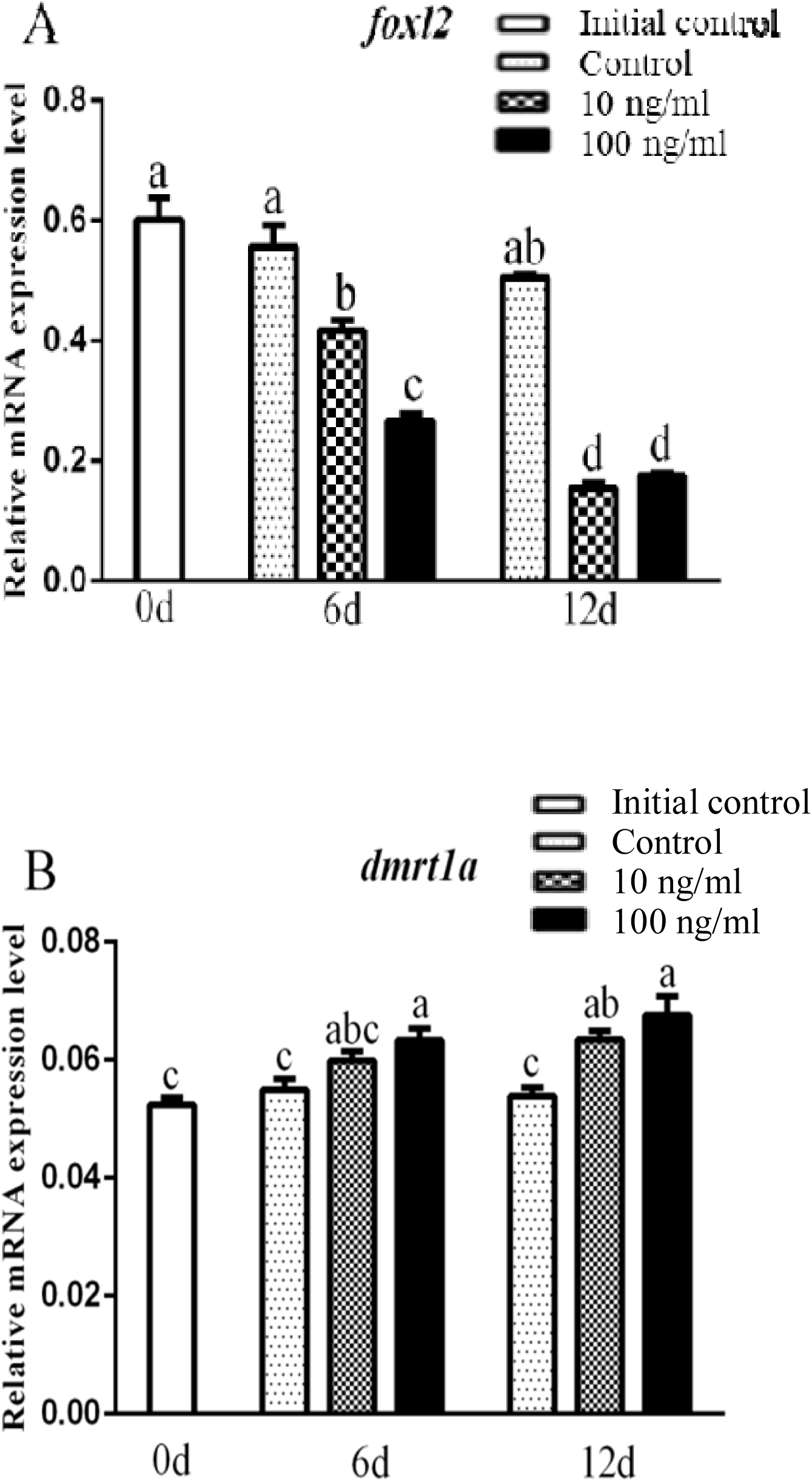
Effects of T on the expression levels of (A) *foxl2* and (B) *dmrt1a* in the ovotestis of *M. albus*. Different letters indicate significant difference between groups within each time (*p* < 0.05).

### Molecular cloning of the full-length 11β-h cDNA

The full length of *11β-h* cDNA sequence was 1812 bp with an open reading frame of 544 amino acids (Supplementary Fig. 2). The amino acid sequence contained without signal peptide cleavage site or transmembrane helix. Several conserved functional motifs were observed including steroid binding site, oxygen-binding region, Ozols’ region, aromatic regions and heme-binding region (Supplementary Fig. 3). We compared the amino acid sequence of *M. albus 11β-h* to that in other species and found 77% identity with *Dicentrarchus labrax*, 76% identity with *Micropogonias undulatus* and 75% identity with *Parajulis poecilepterus* and *Odontesthes bonariensis*. The phylogenetic tree analysis showed that the *11β-h* of *M. albus* and *Epinephelus coioides, P. poecilepterus, D. labrax, M. undulatus, O. bonariensis, O. latipes* and *O. niloticus* were clustered together (Supplementary Fig. 4).

**Fig. 4.**
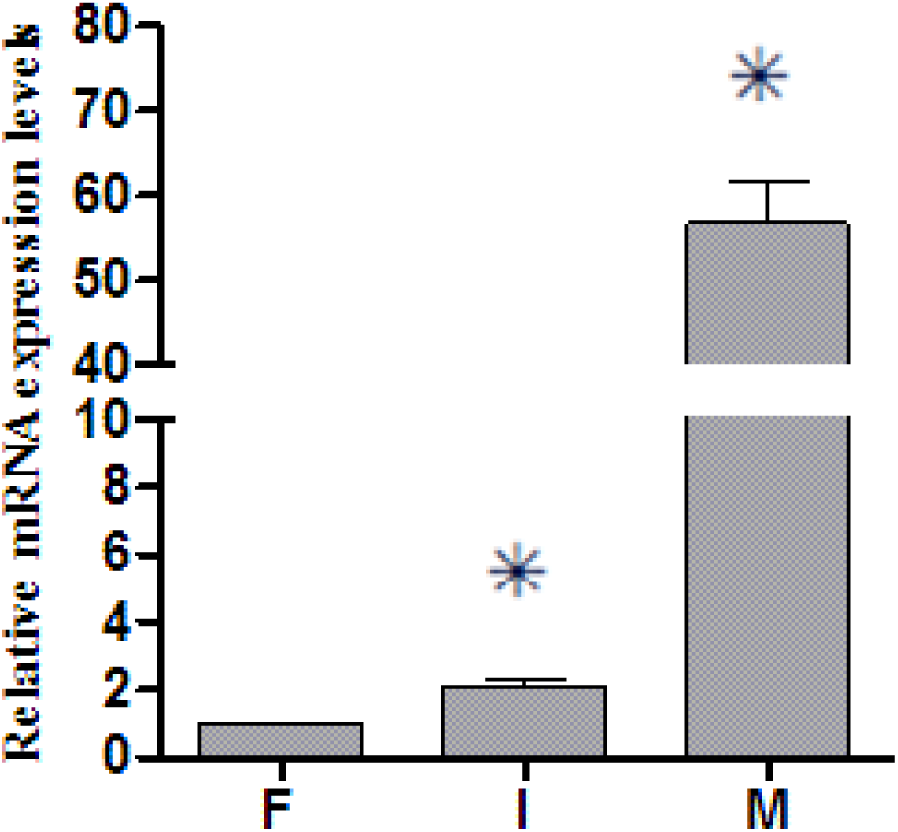
Expression level of *11β-h* during *M. albus* gonadal development. F, ovaries; I, ovotestis; M, testis. (*) indicates significant difference with the former (*p* < 0.05).

### Expression of 11β-H mRNA during sex reversal

*11β-h* was highly expressed in the testis, which was significantly higher than that in the ovary and ovotestis (*p* < 0.05). Moreover, the expression level of *11β-h* in ovotestis was higher than in the ovary (*p* < 0.05) (Fig. 4).

### Molecular cloning of the full-length 11β-hsd2 cDNA

The full length of *11β-hsd2* cDNA sequence was 2267 bp with an open reading frame of 407 amino acids (Supplementary Fig. 5). The amino acid sequence contained without signal peptide cleavage site or transmembrane helix. Several conserved functional motifs were found including NAD-binding domain, *11β-hsd* conserved sequence and catalytic site (Supplementary Fig. 6). We compared the amino acid sequence of *M. albus 11β-hsd2* to that in other species and found 83% identity with *O. latipes*, 80% identity with *O. bonariensis* and 77% identity with *O. niloticus*. The phylogenetic tree analysis showed that the *11β-hsd2* of *M. albus* and *O. bonariensis, O. latipes* and *O. niloticus* were clustered together (Supplementary Fig. 7).

**Fig. 5.**
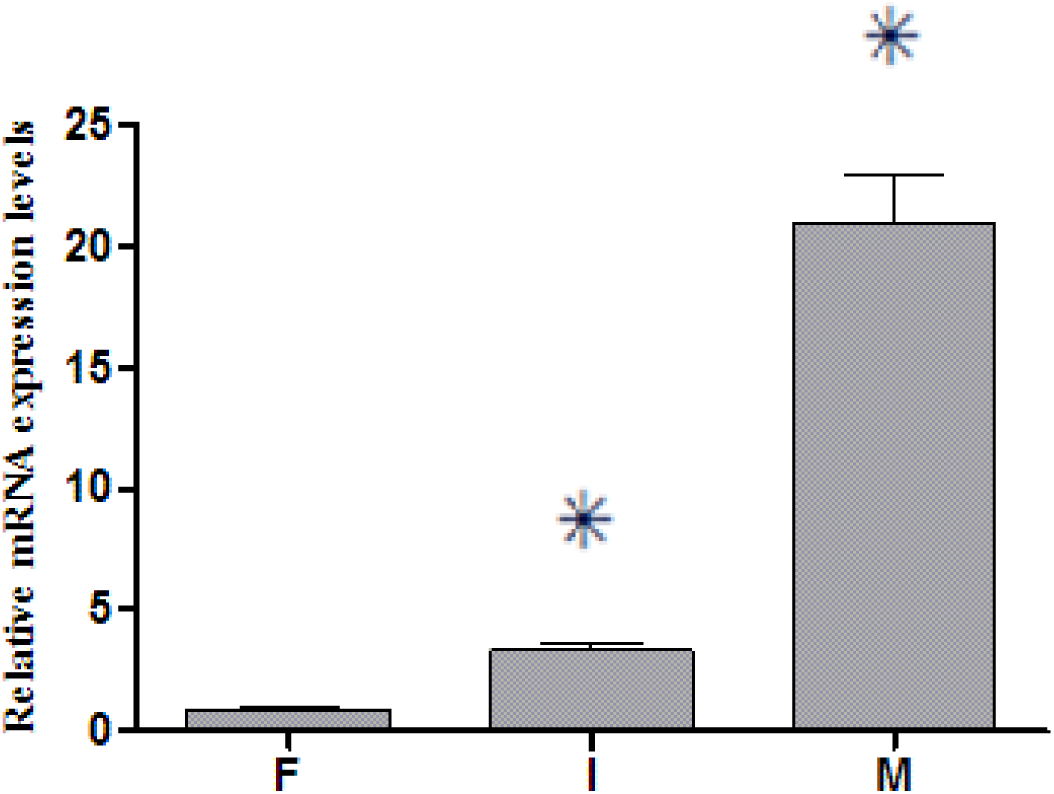
Expression level of *11β-hsd2* during *M. albus* gonadal development. F, ovaries; I, ovotestis; M, testis. (*) indicates significant difference with the former (*p* < 0.05).

**Fig. 6.**
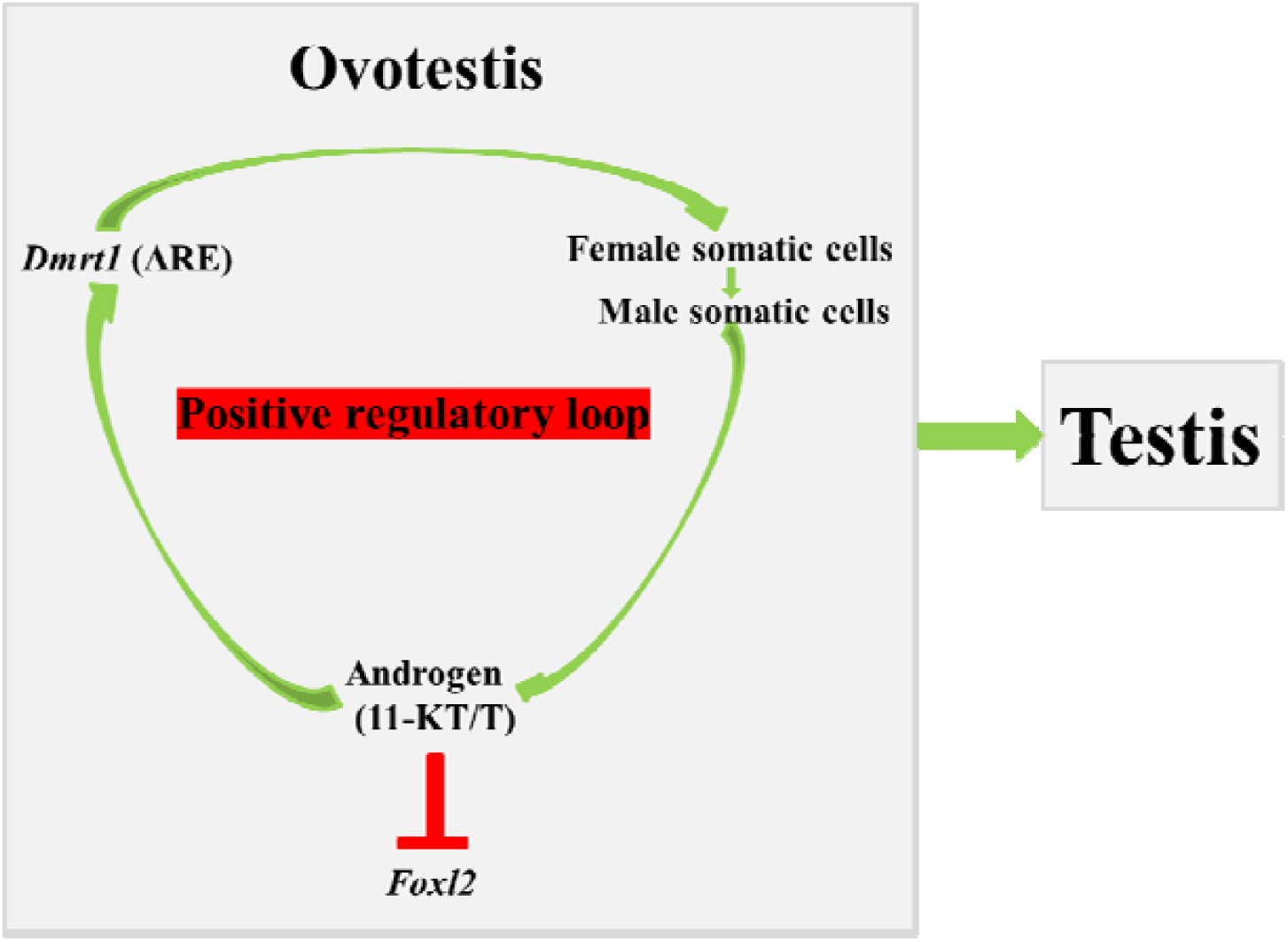
The framework for clarifying the mechanism of *M. albus* sex transdifferentiation. Androgens are synthesized in large amounts in the ovotestis, which activates the transcription of *dmrt1* via putative AREs, resulting in biological effects, which in turn, induces ovarian somatic cells to transdifferentiate into testicular somatic cells. As such, a positive regulatory loop programs the *M. albus* sex reversal. On the other hand, androgens inhibit the expression of *foxl2* and its function.

### Expression of 11β-hsd2 mRNA during sex reversal

*11β-hsd2* was highly expressed in the testis, which was significantly higher than that in the ovary and ovotestis (*p* < 0.05). Moreover, the expression level of *11β-hsd2* in ovotestis was significantly higher than in the ovary (*p* < 0.05) (Fig. 5).

## Discussion

Promoters are, generally, located at the upstream of a transcription start site and have a variety of regulatory motifs, such as the interaction of transcription factors with their corresponding binding sites, which participate in gene regulation (44). In this study, analysis of the promoter region of *dmrt1* showed various transcription binding sites that potentially activated the transcription of *dmrt1*. Specifically, in comparison with the *dmrt1* 5′ upstream region of other known fish species, only in the sequence of *M. albus*, there was one putative ARE on the sense strand (−638 bp ∼ −648 bp), indicating that AR (androgen receptor) was the specific transcription factor of *dmrt1* gene. Sex hormones play an important role in mediating physiological responses and developmental processes through their receptors across all vertebrates. Once androgen ligand binds to AR, the receptor becomes phosphorylated and translocates into the nucleus, in which it binds to ARE(s), and activates the transcription of *dmrt1* gene.

Steroids are known to play a crucial role in gonadal sex differentiation in many non-mammalian vertebrates, but also in the gonadal sex change of hermaphroditic teleosts. *In vitro* culture showed increased expression level of *dmrt1a* but decreased expression level of *foxl2* with increased T concentration and culture time, implying the role of androgen in the transcription of sex-related genes during sex reversal in *M. albus*. Similarly, a hormonal manipulation *in vitro* showed that 11-KT activated the Sertoli cells leading to the completion of spermatogenesis in Japanese eel *Anguilla japonica* (45). Also, Jo et al. (46) observed that the expression levels of *dmrt1* in ovary of *P. olivaceus* were significantly up-regulated by T treatment. Raghuveer et al. (47) observed that methyl testosterone treatment resulted in the initiation of testicular differentiation in juvenile catfish *Clarias gariepinus*, which is supported by specific expression of two forms of *dmrt1*. The expression level of *dmrt1* is high in mature testis of black porgy *Acanthopagrus schlegeli* during sex-reverse process (48). Besides fish species, T-treated ovaries induce upregulated expression of *dmrt1* in the ovotestis of *Rana rugosa* Frogs (49). Aoyama et al. (50) revealed that *dmrt1* was not transcribed at any time during ovarian development but was expressed in the female-to-male sex reversed gonad of amphibians. Hu et al. (35) also observed a high level of *foxl2* expression in the ovary before sex reversal in *M. albus*, but its transcripts decreased sharply when the gonad developed into the ovotestis and testis. Overall, *dmrt1* is essential to maintain vertebrate testis determination (51). *Foxl2* is required to prevent transdifferentiation of an adult ovary to a testis (52). We assumed that the antagonism between *dmrt1* and *foxl2* might cause reprogramming gonad in *M. albus* (53).

We further examined the expression of genes encoding key steroidogenic enzymes during the process of sex reversal in *M. albus*. The expression of gonadal *11β-h* showed obvious sexual dimorphism, with high level in the testis and ovotestis, indicating the vital role of this gene in testis development. Liu et al. (29) also reported that *11β-h* was markedly up-regulated at the onset of testicular development in *M. albus*. Similarly, the expression level of *11β-h* is comparatively low at the early spermatogenesis and sharp increases during spermiogenesis, finally, reaches its highest levels in Atlantic salmon (7). In *O. niloticus*, the expression levels of two isoforms of *11β-h* are detected in testis from 50 days after hatching (dah) onwards and strongly expressed in sex reversed XX testis after fadrozole and tamoxifen treatment, but completely inhibited in 17β-estradiol induced XY ovary (9). In *C. batrachus, 11β-h* is expressed ubiquitously with high levels in testis and could be detected as early as at 0 dah as supported by high level of 11-KT in serum and testicular tissue during pre-spawning and spawning phases, which might facilitate the initiation and normal progression of spermatogenesis (10). The gonadal *11β-hsd2* showed similar expression pattern with *11β-h* in *M. albus*, indicating the vital role of these two genes in the female-to-male reversal. Similarly, *11β-hsd2* is expressed in a wide variety of tissues in *O. niloticus*, with the highest expression in testis (3). Yu et al. (31) found that the expression levels of 17α-hydroxylase, were dominantly expressed in testis, less in ovary, and the least in ovotestis, consistent with the sex reversal process of *M. albus*. Similarly, the expression levels of *11β-h* and *11β-hsd2* are predominantly expressed in testis, much less in ovotestis, and barely in ovary, consistent with a role in the production of 11-KT during sex reversal.

During female-to-male sex reversal, the expression level of *foxl2* is sharply decreased in *M. albus* (35). Also, the aromatase transcripts are decreased when gonad develops into the ovotestis and testis (29). As a result, synthesize of estrogen may decrease during sex reversal. Androgen is the substrate for the production of female hormone, the level of androgen may thus increase. In this study, T also showed a higher inhibitory effect on *foxl2* than positive impact on *dmrt1a*. In this regard, these results are accord with the withdrawal hypothesis of estrogen proposed by Nagahama (54). Moreover, serum T level in female *M. albus* reaches a peak two months after spawning and is significantly higher than the estrogen level (55). Therefore, we suggested that the high level of androgen is the main driving factor for sex reversal in *M. albus*. However, the withdrawal of estrogen during sex reversal is passive, not active, due to the inhibitory action of androgen-*dmrt1a* on the aromatase-*foxl2*.

In conclusion, the gene sequence of *M. albus dmrt1* 5′ upstream region contained two unique AREs, indicating that AR was the specific transcription factor of *dmrt1*. Also, the *dmrt1a* was positive regulated by T, suggesting that the blood androgen could promote the transcription of *dmrt1* during sex reversal. Moreover, high expression levels of *11β-h* and *11β-hsd2* were observed during female-to-male sex reversal, indicating the large production of 11-KT during this process. Overall, as shown in Fig. 6, androgens are synthesized in large amounts in *M. albus* during sex reversal, promoting the transcription of *dmrt1* via putative ARE(s), which in turn, induces ovarian somatic cells to transdifferentiate into testicular somatic cells.

## Methods

### Fish

The wild *M. albus* (body weight ∼200 g) were collected from Hubei, China and transported to the Fish Breeding Laboratory, Shanghai Ocean University (Shanghai, China). After 30 days of acclimation, the animals were sacrificed by anesthesia with MS-222 and dissected on ice. A portion of the gonad was fixed in Bouin’s fluid for histological assessment of the sexual status. The other samples were frozen in liquid nitrogen and stored at −80°C. All experiments were performed with the approval from the Institutional Animal Care and Use Committee of Shanghai Ocean University.

### Isolation of 5’ upstream region of dmrt1 and sequence analysis

Genomic DNA was isolated from gonad tissue by using manufacturer’s protocol (Qiagen, GmbH, Germany). The integrity of DNA was checked using 2% agarose gel electrophoresis. Based on the DNA sequence of *M. albus dmrt1* obtained from NCBI (Accession No: NW-018128265), the specific primers (Supplementary Table 1) were designed to amplify the 5’ upstream region of *dmrt1* gene. The JASPAR database and associated tools (http://jaspar.genereg.net), Match (BioBase), AliBaba2.1 (Biobase) and MOTIF (GenomeNet) were used to predict the transcription factor binding sites (56-58).

### Histology and light microscopy observation

The dissected gonads were stored in 4% paraformaldehyde for 24□h. After rinsing with flowing water, the gonads were dehydrated in a series of ethanol, embedded in paraffin and cut by a microtome at 6 μm thickness. After hematoxylin-eosin dye, the stained sections were observed under an inverted phase-contrast microscope (Olympus BX-53, Tokyo, Japan).

### In vitro culture

The ovotestis (Supplementary Fig. 1) was cut into 1 × 1 × 0.5 mm^3^ small pieces, washed three times with PBS × 1, and then transferred to 24-well culture plates. The control group was cultured in a medium containing 15% fetal bovine serum and 1% penicillin/streptomycin. The treatment groups were cultured in a medium containing additional 10 (low) or 100 ng/ml (high) of T. The gonadal tissues were cultured in CO_2_ incubator at 27 □. One half of the medium was changed every other day. The growth of cell was observed under an inverted microscope daily and sampled on day 6 and day 12 for *dmrt1a* and *folx2* expression analysis.

### Total RNA extraction and cDNA synthesis

Total RNA was extracted using Trizol method (Invitrogen, USA) according to the manufacturer’s instructions. The quality of total RNA was determined by using a NanoDrop 2000 spectrophotometer (Thermo Fisher Scientific, USA) measured at 260/280□nm, and the integrity was screened by 1.5% agarose gel electrophoresis. The cDNA was synthesized by using a PrimeScript™ RT reagent Kit (Takara, China) following the manufacturer’s instructions. The obtained cDNA templates were stored at −80□°C for gene cloning and qRT-PCR amplification.

### Cloning the full-length cDNA of 11β-h and 11β-hsd2 gene and sequence analysis

The primers (Supplementary Table 1) were designed to amplify the internal region of *11β-h* and *11β-hsd2* gene respectively by using Takara PCR Amplification Kit (Takara, Japan). To obtain the full-length cDNA sequences, 3′ and 5′ rapid-amplification of cDNA ends Polymerase Chain Reaction (RACE-PCR) was carried out by using the SMART™ RACE cDNA Amplification Kit (Clontech, USA) according to the manufacturer’s instructions. The amplified PCR products were excised by 1.5% agarose gel electrophoresis, and bands of expected size were dissociated and purified by using a gel extraction kit (Omega, China). The PCR products were directly ligated into PMD19-T simple vector (TaKaRa, China) and then transformed into Escherichia coli BL21 competent cells (Transgen, China).

Prediction of the open reading frame on *11β-h* and *11β-hsd2* was performed by using the BLAST Program of NCBI (http://www.ncbi.nlm.nih.gov/blast). Prediction of the protein domains were carried out by using the SMART program (http://www.smart.emble-heidelberg.de/), InterPro (http://www.ebi.ac.uk/interpro/) and IMGT (http://www.imgt.org/). Prediction of signal peptide was performed by using the SignalP 4.0 (http://www.cbs.dtu.dk/services/SignalP/). Conserved motifs were identified by using Conserved Domain Search Service from NCBI. Multiple alignments of amino acid sequences were performed by using the ESPript (http://multalin.toulouse.inra.fr/multalin/). The phylogenetic neighbor-joining (NJ) tree was constructed by using the MEGA 6.0 program (59), and the reliability was assessed by 1000 bootstrap replications.

### Quantitative real-time PCR and expression analysis

The expression levels of *dmrt1a, foxl2, 11β-h* and *11β-hsd2* were quantified by real-time quantitative RT-PCR with specific primers (Supplementary Table 1), by using SYBR Premix Ex *Taq* II (Tli RNaseH Plus) Kit (Takara, Dalian, China) and CFX96TM real-time system (Bio-Rad, Hercules, CA, USA). At the end of the reactions, the credibility of the qRT-PCR was analyzed through melting curve. All samples were run in triplicate, and each assay was repeated three times. After finishing the program, the cycle threshold (Ct) value was automatically determined by the Bio-Rad CFX Manager software. The mRNA expression levels were calculated relative to β-actin using the 2^−ΔΔCt^ method (60).

### Statistical analysis

Raw data were assessed for the normality of distribution and the homogeneity of variance with the Kolmogorov-Smirnov test and Levene’s test, respectively. The data conformed to a normal distribution and were suitable for testing with analysis of variance (ANOVA). The differences in mRNA expression levels of *dmrt1a, foxl2, 11β-h* and *11β-hsd2* between treatments were compared with one-way ANOVA at the significance level of 0.05 (*p* < 0.05). Data analyses were performed using the software SPSS for Windows (Release 20.0).

## Acknowledgements

This research is funded by Ministry of Agriculture, Science and Technology Education Department (201003076).

## Author contributions

B.W. and X.Q. designed the research and drafted the paper, L.P. conducted the cell culture, J.G. isolated of 5’ upstream region of *dmrt1*, H.W. conducted the gene cloning, Q.W. conducted the gene expression.

