## Supplemental Tables and Figures for "Androgen-*dmrt1* positive feedback programs the rice field eel (*Monopterus albus*) sex transdifferentiation"

**Fig. S1.** Microscopic examination on gonad tissue during stage I ovotestis in *M. albus*. II: stage II oocyte; IV: stage IV oocyte; GF: genital fold.

**Fig. S2.** Nucleotide and deduced amino acid sequences of *11β-h*. The 5′ and 3′ untranslated regions are in lower cases. The translation of stop codon is designated with an asterisk.

**Fig. S3.** Alignment of the amino acid sequences of *11β-h* gene in *M. albus* with other species. Rectangles indicate conserved domains. Mo: *Monopterus albus*; Dic: *Dicentrarchus labrax* (AF449173); Mic: *Micropogonias undulates* (EU673091); Odo: *Odontesthes bonariensis* (GQ381267); Onc: *Oncorhynchus mykiss* (AF179894); Dan: *Danio rerio* (DQ650710); Ore: *Oreochromis niloticus* (ACY39528); Ory: *Oryzias latipes* (NM_001105100); Hom1: *Homo sapiens1* (NM_000497); Hom2: *Homo sapiens2* (P19099); Rat: *Rattus norvegicus1* (NM_012537); Ran: *Rana catesbeiana* (D10984).

**Fig. S4.** Phylogenetic tree of *M. albus* and other species *11β-h* amino acid sequence. Mon: *Monopterus albus*; Dic: *Dicentrarchus labrax* (AF449173); Mic: *Micropogonias undulatus* (EU673091); Epi: *Epinephelus coioides* (JQ320138); Odo: *Odontesthes bonariensis* (GQ381267); Par: *Parajulis poecilepterus* (AB911711); Onc1: *Oncorhynchus mykiss* (AF179894); Onc2: *Oncorhynchus mykiss* (AF217273); Ore1: *Oreochromis niloticus* (FJ713103); Ore2: *Oreochromis niloticus* (FJ713104); Dan: *Danio rerio* (DQ650710); Ory: *Oryzias latipes* (NM_001105100); Ovi: *Ovis aries* (OAU78478); Hom1: *Homo sapiens1* (NM_000497); Hom2: *Homo sapiens2* (P19099); Rat1: *Rattus norvegicus1* (NM_012537); Rat2: *Rattus norvegicus2* (P30099); Rat3: *Rattus norvegicus3* (P30100); Ran: *Rana catesbeiana* (D10984).

**Fig. S5.** Nucleotide and deduced amino acid sequences of *11β-hsd2.* The 5′ and 3′ untranslated regions are in lower cases. The translation of stop codon is designated with an asterisk.

**Fig. S6.** Alignment of the amino acid sequences of *11β-hsd2* gene in *M. albus* with other species. Rectangles indicate conserved domains. Mon: *Monopterus albus*; Gob: *Gobiocypris rarus* (KC454276); Dan: *Danio rerio* (NM_212720); Cla: *Clarias gariepinus* (GU220074); Onc: *Oncorhynchus mykiss* (NM_001124218); Ang: *Anguilla japonica* (AB252646); Ore: *Oreochromis niloticus* (NM_001279757); Odo: *Odontesthes bonariensis* (ADR30382); Hom: *Homo sapiens* (BC036780); Mus: *Mus musculus* (BC066209); Sus: *Sus scrofa* (NM213913).

**Fig. S7.** Phylogenetic tree of *M. albus* and other species *11β-hsd2* amino acid sequence. Mon: *Monopterus albus*; Gob: *Gobiocypris rarus* (KC454276); Dan: *Danio rerio* (NM_212720); Cla: *Clarias gariepinus* (GU220074); Onc: *Oncorhynchus mykiss* (NM_001124218); Ang: *Anguilla japonica* (AB252646); Ore: *Oreochromis niloticus* (NM_001279757); Odo: *Odontesthes bonariensis* (ADR30382); Ory: *Oryzias latipes* (ABK59971); Hom: *Homo sapiens* (BC036780); Mus: *Mus musculus* (BC066209); Sus: *Sus scrofa* (NM213913).

**Table S1.** List of primers used for cloning, qPCR and promoter motif analysis.

**Table S2.** Sequence alignment of the 5′ flanking region of *dmrt1* for *M. albus* and other species.

Fig. S1

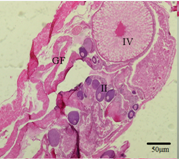

Fig. S2

1 gtttgcact	ATG AGG AGC ATG TAC ATG GGG GTG ACA GTG TGT GTC	45

1	M	R	S	M	Y	M	G	V	T	V	C	V	12

46 AGA GCA CAG GGC ACC TGT GGC TCC ACA AAT CTG CTT TGT GGT GTC	90

13	R	A	Q	G	T	C	G	S	T	N	L	L	C	G	V	27

91 ATG CTG CAA AGA GCT TTT TGT ATG ACT GCA GCA GAG ACA GTG GCA	135

28	M	L	Q	R	A	F	C	M	T	A	A	E	T	V	A	42

136 AAG TCA GAG AGG AGT AAA GGA GTT AGA AAA AGG GGT GTG GAT GGA	180

43	K	S	E	R	S	K	G	V	R	K	R	G	V	D	G	57

181 CGA GTA CGG CGC TTT GAG GAG ATC CCT CAC ACG GGC AGG AGT GGT	225

58	R	V	R	R	F	E	E	I	P	H	T	G	R	S	G	72

226 TGG ATC AAC CTG GTG AAG TTC TGG AGA GAA GGC CGT TTT GGG CAA	270

73	W	I	N	L	V	K	F	W	R	E	G	R	F	G	Q	87

271 ATT CAC AAA CAT ATG GGG CGC ACC TTC AGC GCC CTT GGC CCC ATC	315

88	I	H	K	H	M	G	R	T	F	S	A	L	G	P	I	102

316 TAC AGG GAG CAC GTG GCC ACG CAA AGC AGT GTG AAC ATC TTT CTT	360

103	Y	R	E	H	V	A	T	Q	S	S	V	N	I	F	L	117

361 CCA TCC GAC ATC GGT GAG CTG TTT CGC TCT GAG GGC CTG TAC CCT	405

118	P	S	D	I	G	E	L	F	R	S	E	G	L	Y	P	132

406 GAA CGG ATG ACC GTG CAG CCT TGG GTG ACG CAC AGG GAG ATA CGC	450

133	E	R	M	T	V	Q	P	W	V	T	H	R	E	I	R	147

451 CAT CAC AGC AAG GGA GTC TTC CTC AAG AAT GGT GAG GAG TGG CGA	495

148	H	H	S	K	G	V	F	L	K	N	G	E	E	W	R	162

496 GCT GAC CGT CTA CTG CTC AAC AAA GAG GTG ATG ATG AAT GTG GCC	540

163	A	D	R	L	L	L	N	K	E	V	M	M	N	V	A	177

541 GTG CAG CGC TTT CTC CCT CAC CTT GAT GAG GTG GCG AGG GAC TTC	585

178	V	Q	R	F	L	P	H	L	D	E	V	A	R	D	F	192

586 TGT CGA ATG CTG CAG GCG AGA GTG GAG AAG GAG GGA AGA GTC GTG	630

193	C	R	M	L	Q	A	R	V	E	K	E	G	R	V	V	207

631 GAA GGG AAA CAA AGT CTG ACC ATT GAA CCC AGT CCT GAC CTC TTC	675

208	E	G	K	Q	S	L	T	I	E	P	S	P	D	L	F	222

676 CGC TTT GGG CTG GAA GCC ATT TGC CAT GTG CTC TAT GGG GAG CGC	720

223	R	F	G	L	E	A	I	C	H	V	L	Y	G	E	R	237

721 ATG GGC CTC TTC TCC TCA TCT CCC TCC ATG GAG TCT CAG AAG TTC	765

238	M	G	L	F	S	S	S	P	S	M	E	S	Q	K	F	252

766 ATC TGG GCT TTG AAG CAG ATG CTG GCA ACC ACC AAT CCT CTT CTT	810

253	I	W	A	L	K	Q	M	L	A	T	T	N	P	L	L	267

811 TAC CTG CCC CAT CGC CTG CTG CTC CGC GTC GGT GCT CCG CTG TGG	855

268	Y	L	P	H	R	L	L	L	R	V	G	A	P	L	W	282

856 GTC CAG CAC GCC AGT GCA TGG GAC TAC ATC TTC TCT CAT GCG GAT	900

283	V	Q	H	A	S	A	W	D	Y	I	F	S	H	A	D	297

901 GTG AGG ATC CAG AGG GGA TAC CAG CGC CAG TCA TCC TCC CGG GGT	945

298	V	R	I	Q	R	G	Y	Q	R	Q	S	S	S	R	G	312

946 CGA AGG TCT GAG GCT GGA GCA GCT GGA GAC CGC CAC ACT GGG GTT	990

313	R	R	S	E	A	G	A	A	G	D	R	H	T	G	V	327

991 CTG GGC CAA CTC CTA GAG AAA GGA CAA CTA TCT ATG GAC CTG ATC 1035

328	L	G	Q	L	L	E	K	G	Q	L	S	M	D	L	I	342

1036 AAA GCC AAC ATG ACT GAG CTG ATG GTT GGA GGG GTC GAC ACG ACA 1080

343	K	A	N	M	T	E	L	M	V	G	G	V	D	T	T	357

1081 GCT GTG CCC CTG CAG TTT GCT CTG TTT GAG CTG GCC CGC AAC CCA 1125

358	A	V	P	L	Q	F	A	L	F	E	L	A	R	N	P	372

1126 GAG GTG CAG GAG AAG GTG AGG CAG CAG GTG ATC GAG TCT TGG GCG 1170

373	E	V	Q	E	K	V	R	Q	Q	V	I	E	S	W	A	387

1171 CAG GCG GCT GGT GAC CTT CAG AAA GCT CTG CAG GGG GTG CCA CTG 1215

388	Q	A	A	G	D	L	Q	K	A	L	Q	G	V	P	L	402

1216 CTA AAA GGC ACT GTC AAA GAG ACC CTT AGG TTA TAC CCG GTG GCA 1260

403	L	K	G	T	V	K	E	T	L	R	L	Y	P	V	A	417

1261 ATC ACA GTA CAA CGG CGT CCA GTC AAA GAT GTT GTT CTT CAG AAT 1305

418	I	T	V	Q	R	R	P	V	K	D	V	V	L	Q	N	432

1306 TAC CAC ATA CCA GCA GGG ACA ATG GTC CAG GTC TGT CTT TAC CCC 1350

433	Y	H	I	P	A	G	T	M	V	Q	V	C	L	Y	P	447

1351 CTG GGA AGG AGT GCA GAG GTG TTT GAC AAT CCA GAG CGC TTT GAC 1395

448	L	G	R	S	A	E	V	F	D	N	P	E	R	F	D	462

1396 CCT GAG CGG TGG TGC AGC ATT AGA GAA GGT GGC CAG AGA CCG GAG 1440

463	P	E	R	W	C	S	I	R	E	G	G	Q	R	P	E	477

1441 TTG GGG TTT CGC TTC ATG GCT TTC GGT TTA GGG GCA AGG CAA TGT 1485

478	L	G	F	R	F	M	A	F	G	L	G	A	R	Q	C	492

1486 GTC GGA CGG AGG ATT GCT GAG AAT CAG ATG CAG CTC TTA CTC ATG 1530

493	V	G	R	R	I	A	E	N	Q	M	Q	L	L	L	M	507

1531 CGT GTC CTG CTG AGC TTC CAT CTC AGC GTG CAG TCC TCA GAG GAC 1575

508	R	V	L	L	S	F	H	L	S	V	Q	S	S	E	D	522

1576 GTC AAG ACC ACA TTC GCC TTC CTC CTC CAG CCT GAG ACT CCG CCG 1620

523	V	K	T	T	F	A	F	L	L	Q	P	E	T	P	P	537

1621 AGG ATC ACG TTC AGC CGG ATC TGA	1644

538	R	I	T	F	S	R	I	*	544

1645 cgcacacagagaagagggttgtcagcatttgacatgtttaatcacacaatttagcc	1700

1701 tgtgtcatcttaacaacactgtcataaactccgtaatcacgccaagaatttgtaat	1756

1757 gctgtcaaataaagtatcatttcctcctccaaaaaaaaaaaaaaaaaaaaaaaaaa	1812

Fig. S3

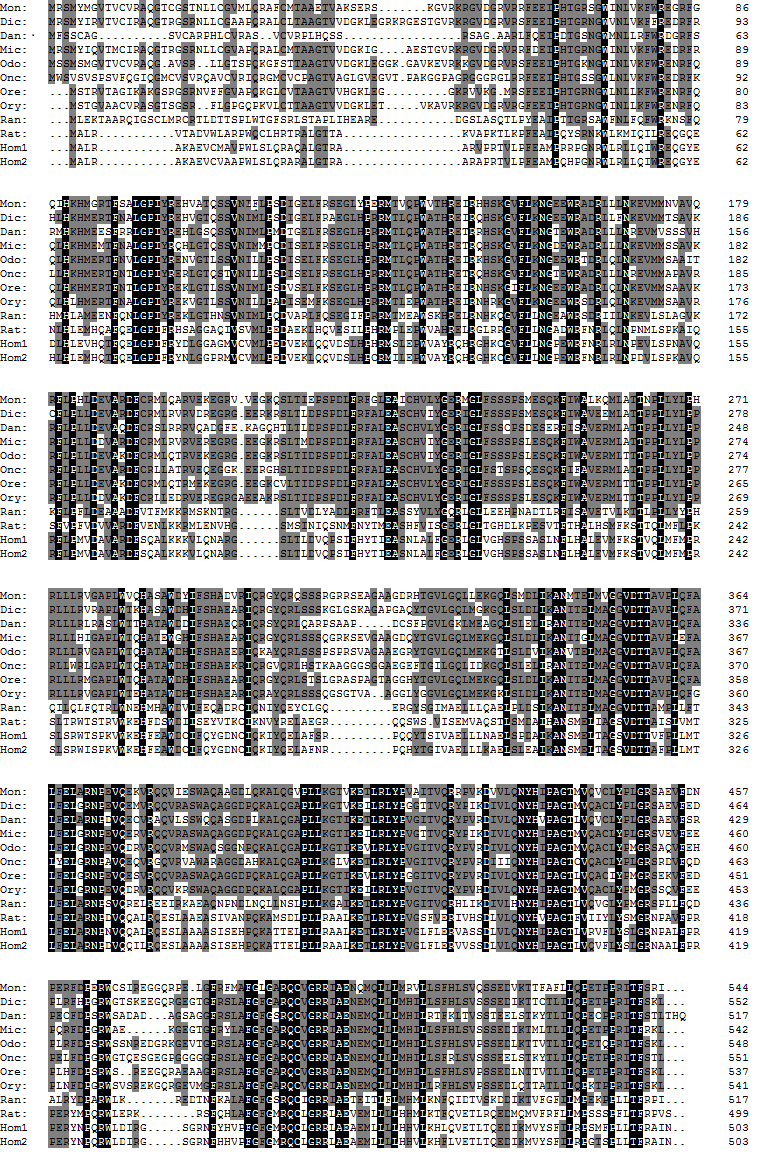

Fig. S4

Epi|JQ320138

Par|AB911711

Dic|AF449173

Mic|EU673091

Odo|GQ381267

Ory|NM_001105100

Ore1|FJ713103

Ore2|FJ713104

Mon

Onc1|AF179894

Onc2|AF217273

Dan|DQ650710

Ran|D10984

Rat1|NM_012537

Rat3|P30100

Rat2|P30099

Ovi|OAU78478

Hom1|NM_000497

Hom2|P19099

98

100

100

89

100

100

100

100

68

98

85

85

51

58

51

77

0.1

Fig. S5

1 acatggggagctctgcactggctgcactgaaccgaccagacacagtgtgtgcagcaca	58

59 gagggagagcgatcaagagacataaacactgactgacactaacctggatcagactcgt	116

117 ttcttgttcttgtttaaatctgaaggtgcagaggcacagacagagaacactcagacaa	174

175 gtaaaccaggtgaacctaatg	195

196 ATG GAC GAC TAC ACC CTC CCC TTC TGG ATC TAC CTG TGT GTT CTT	240

1	M	D	D	Y	T	L	P	F	W	I	Y	L	C	V	L	15

241 ACT GTG TTT GTT GGT GGT GCT ATG AAG ATG ATT TTG GCG TCC CAC	285

16	T	V	F	V	G	G	A	M	K	M	I	L	A	S	H	30

286 CTG AGC ACC GCC TCT ACC CTG GTG GCC TGG CTG GGA GCC ACT GTG	330

31	L	S	T	A	S	T	L	V	A	W	L	G	A	T	V	45

331 CTG GTG GAG CGT CTG TGG GCG TTC TGT CTG CCG GCC ATG CTG CTG	375

46	L	V	E	R	L	W	A	F	C	L	P	A	M	L	L	60

376 CTG GTG CTG CTT GGC CTC ACC TGT TGC TTC TAC AAT GCC ACC AGG	420

61	L	V	L	L	G	L	T	C	C	F	Y	N	A	T	R	75

421 AGT GCC CCC CTG CCC ACC ACC CTT CCT GCC CAT GGC AAG GCT GTC	465

76	S	A	P	L	P	T	T	L	P	A	H	G	K	A	V	90

466 TTC ATC ACA GGC TGT GAC TCT GGT TTT GGA AAT GCC ACA GCA AAG	510

91	F	I	T	G	C	D	S	G	F	G	N	A	T	A	K	105

511 CGT CTG GAC GCC ATG GGC TTT GAG GTG TTT GCC ACG GTT TTG GAC	555

106	R	L	D	A	M	G	F	E	V	F	A	T	V	L	D	120

556 CTG TCC GGA GAT GGA GCC AGG GAG CTG CAG AGG ACC TGC TCC CCC	600

121	L	S	G	D	G	A	R	E	L	Q	R	T	C	S	P	135

601 CGC CTC ACC CTC CTC CAG GTG GAC ATC ACT CAG CCG CAG CAG ATC	645

136	R	L	T	L	L	Q	V	D	I	T	Q	P	Q	Q	I	150

646 CAG CAG GCG CTG CTG GAC ACC AGG ACC AAA CTG GGC CTC AAA GGT	690

151	Q	Q	A	L	L	D	T	R	T	K	L	G	L	K	G	165

691 TTG TGG GGT TTG GTA AAC AAT GCT GGA CGG TGT GTG AAC ATC GGA	735

166	L	W	G	L	V	N	N	A	G	R	C	V	N	I	G	180

736 GAT GCC GAG CTG TCG CTG ATG TCC AAC TTC CGC GGC TGC ATG GAG	780

181	D	A	E	L	S	L	M	S	N	F	R	G	C	M	E	195

781 GTC AAC TTC TTC GGC ACA CTG AGC GTC ACC AAG TGC TTC CTG CCA	825

196	V	N	F	F	G	T	L	S	V	T	K	C	F	L	P	210

826 CTG CTG CGT CAG GCC AAA GGA CGG ATC GTC ACC ATC TCC AGC CCT	870

211	L	L	R	Q	A	K	G	R	I	V	T	I	S	S	P	225

871 GCT GGT GAC CAC CCG TTT CCC TGT CTG GCA GCG TAC GGA GCA TCT	915

226	A	G	D	H	P	F	P	C	L	A	A	Y	G	A	S	240

916 AAA GCA GCT CTC AAC CTC TTC ATC AAC ACT CTG CGG CAC GAG CTC	960

241	K	A	A	L	N	L	F	I	N	T	L	R	H	E	L	255

961 GAG CCC TGG GGT GTG CAA GTT TCA ATC ATC CTG CCA TCC GCG TTC	1005

256	E	P	W	G	V	Q	V	S	I	I	L	P	S	A	F	270

1006 AAG ACA GGT CAT CCC AGT AAC TAT GCG TAC TGG GAG CAG CAG CAC	1050

271	K	T	G	H	P	S	N	Y	A	Y	W	E	Q	Q	H	285

1051 AAA CAG CTG CTG CAG AAT CTG TCT CCG GCG CTG CTC GAA GAC TAC	1095

286	K	Q	L	L	Q	N	L	S	P	A	L	L	E	D	Y	300

1096 GGT GAG GAC TAT GTG ACC GAG ACC AAG GAC CTC TTC CAT AGC CAT	1140

301	G	E	D	Y	V	T	E	T	K	D	L	F	H	S	H	315

1141 GCC AGC CAG GCC AAC CCT GAC CTC AGC CCT GTT GTG GAC GCC ATC	1185

316	A	S	Q	A	N	P	D	L	S	P	V	V	D	A	I	330

1186 ATC CAT GCG CTG CTG GCG CCA CAG CCG CAG GCA CGT TAC TTT GCG	1230

331	I	H	A	L	L	A	P	Q	P	Q	A	R	Y	F	A	345

1231 GGG CCT GGT GTC GGC CTC ATG TAC TTC ATC CAG ACC TAC TGC CCC	1275

346	G	P	G	V	G	L	M	Y	F	I	Q	T	Y	C	P	360

1276 TTC AGT GTC AGC AAC CGC TTC CTC CAG AAA ATG TTT ATG AAG AAG	1320

361	F	S	V	S	N	R	F	L	Q	K	M	F	M	K	K	375

1321 AAG CTG ACA CCT TAC GCC CTG AGG AAG CAA TCG GGC TTC AAC CTG	1365

376	K	L	T	P	Y	A	L	R	K	Q	S	G	F	N	L	390

1366 AAC CTC AGC CTC CAC AAC AAC AAT AAC AAC AAC AAC GGG GAA AAA	1410

391	N	L	S	L	H	N	N	N	N	N	N	N	G	E	K	405

1411 CTC ATG TAG	1419

406	L	M	*	407

1420 ctatgacacagtagacacatcctgatggtatcaaaagcagcgatagaggaccacttct	1477

1478 gtaacacactcatagagctgtaacttgtccttttctatgtgcagccatgggtcaatca	1535

1536 gtgaaaccgtcacagtcagagagatcagctacactgtactgagccacactcactgcac	1593

1594 gtgccttcatctcacaaaaggtcacagagagaatccagcactacagcagcttgtttaa	1651

1652 atactgactccaagattacttaaaaacagatacaaacatccgtgcacttgtttgccag	1709

1710 ggtggacgcactcagcctcgctgtgggcctgttacagctgtcatcttcagtgttgtca	1767

1768 ggagacagccatgtcatcagcctgccaacatcacaccccagctagccagtagagcccg	1825

1826 gctccgtctcaggcccgacccagctgctgctgcttggaacttactttcagccacgttt	1883

1883 cacatgcacacagtgaaccaaatctggtgggaatgtctgggccggagacagctcacag	1941

1942 ccaactgtgtgcatacaattggcctgactccgctggatccctgagatggtgtgaagcc	1999

2000 aaccatgaatgttaaactgtgagacgttcacccagttaccaatgccagtgcttctatc	2057

2058 aaacagtcagacttccatcccaaggtttattgtacgacggcggacgtcgatgcagtgt	2115

2116 cggggaagctactctgatagtgtagctttacaagctaccaattgagctgcagcaaatc	2173

2174 tactgtaaggagagattgagtctggttaagtaaagctacttagaaaaggcagtttaat	2231

2232 ctactttgtataaaaaaaaaaaaaaaaaaaaaaaaa	2267

Fig. S6

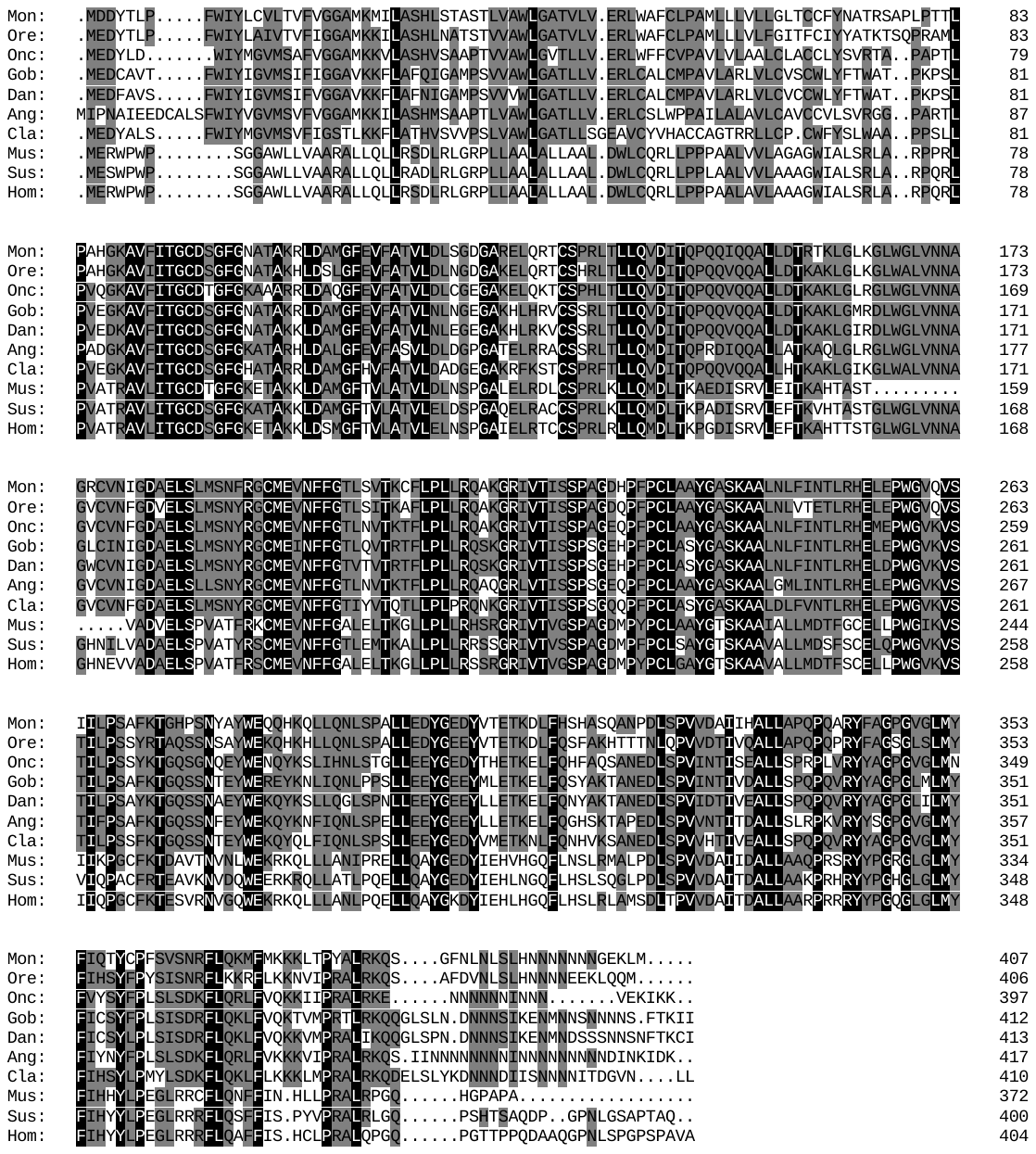

Fig. S7

Gob|KC454276

Dan|NM_212720

Cla|GU220074

Onc|NM_001124218

Ang|AB252646

Mon

Ore|NM_001279757

Odo|ADR30382

Ory|ABK5997146

Sus|NM_213913

Mus|BC066209

Hom|BC036780

83

96

98

56

75

100

75

37

41

0.05

Table S1

| Primer | Nucleotide sequence (5'-3') |
| --- | --- |
| *dmrt1-*F1 (Walking) | CAAAAGGGAGCGTTTCAGATAC |
| *dmrt1-*R1 (Walking) | GGGGACAGAGGTAGGGTGC |
| *dmrt1-*F2 (Walking) | ATTAGAATATCGTGGACAA |
| *dmrt1-*R2 (Walking) | ACAACTATGCTGCCAACT |
| *dmrt1a*-F (real time) | GATGTCAAGGCTGAGTGCGA |
| *dmrt1a*-R (real time)  *foxl2*-F (real time)  *foxl2*-R (real time) | CCACGAGGCTAAGAAGGAACT  TGACAACAACACGAAGAAGGAG  GGCAATGAGAGCAACATAGGA |
| *18S rRNA*-F | GTGGAGCGATTTGTCTGGTTA |
| *18S rRNA*-R | CGGACATCTAAGGGCATCAC |
| 11β-h-F1  11β-h-R1  11β-hsd2-F1  11β-hsd2-R1 | AGYCCTGACCTCTTCCGCTT GTCCCWGCWGGTATGTGGTA  GTCTTCATCACAGGYTGTG  GGAASAGHTCCTTGGTCTC |
| RACE-*11β-h*-F1  RACE-*11β-h*-F2  RACE-*11β-h*-R1  RACE-*11β-h*-R2  RACE-*11β-hsd2*-F1  RACE-*11β-hsd2*-F2  RACE-*11β-hsd2*-R1  RACE-*11β-hsd2*-R2 | GACTCGATCACCTGCTGCCTCACCTTCT  CCAGCCCAAAGCGGAAGAGGTCAGGACT  GCAGGAGAAGGTGAGGCAGCAGGTGATCG  CGGTGGCAATCACAGTACAACGGCGTCC  GCCGAAGAAGTTGACCTCCATGCAGCC  GCCCATGGCGTCCAGACGCTTTGCTGTG  CGCGGCTGCATGGAGGTCAACTTCTTCGG  GGTGAGGACTATGTGACCGAGACCAAGG |
| 11β-h-F2 (real time)  11β-h-R2 (real time)  11β-hsd2-F2 (real time)  11β-hsd2-R2 (real time) | TCCTGACCTCTTCCGCTTTGG CCTCTGGATCCTCACATCCGC  GCTCTCAACCTCTTCATCAACAC CATAGTCCTCACCGTAGTCTTCG |
| β-actin F  β-actin R | ATCGCCGCACTCGTTGTTGAC  CCTGTTGGCTTTGGGGTTC |

Table S2

| Fish species | Full length (bp) | Number | Position | Start (bp) | End (bp) |
| --- | --- | --- | --- | --- | --- |
| Rice-field eel *Monopterus albus* | 1421 | 2 | Sense strand | -638 | -648 |
|  |  |  | Antisense strand | -903 | -917 |
| Chinese wrasse *Halichoeres tennispinis* | 1537 | 1 | Antisense strand | -821 | -835 |
| Atlantic cod  *Gadus morhua* | 1175 | 1 | Antisense strand | -463 | -477 |
| Zebrafish  *Danio rerio* | 1500 | 0 |  |  |  |
| Barramundi perch  *Lates calcarifer* | 1500 | 0 |  |  |  |
| Common carp  *Cyprinus carpio* | 1500 | 0 |  |  |  |
| Japanese medaka  *Oryzias latipes* | 1500 | 0 |  |  |  |
